# Evolution within the fungal genus *Verticillium* is characterized by chromosomal rearrangement and gene loss

**DOI:** 10.1101/164665

**Authors:** Xiaoqian Shi-Kunne, Luigi Faino, Grardy C.M. van den Berg, Bart P.H.J. Thomma, Michael F. Seidl

**Affiliations:** Laboratory of Phytopathology, Wageningen University, Droevendaalsesteeg 1, 6708 PB Wageningen, The Netherlands

## Abstract

The fungal genus *Verticillium* contains ten species, some of which are notorious plant pathogens causing vascular wilt diseases in host plants, while others are known as saprophytes and opportunistic plant pathogens. Whereas the genome of *V. dahliae*, the most notorious plan pathogen of the genus, has been well characterized, evolution and speciation of other members of the genus received little attention thus far. Here, we sequenced the genomes of the nine haploid *Verticillium* spp. to study evolutionary trajectories of their divergence from a last common ancestor. Frequent occurrence of chromosomal rearrangement and gene family loss was identified. In addition to ~11,000 core genes that are shared among all species, only 200-600 species-specific genes occur. Intriguingly, these species-specific genes show different features than core genes.

## INTRODUCTION

Species continuously evolve by genetic variation that enables adaptation to changing and novel environments. In many eukaryotes, such genomic variation is established during sexual reproduction where genetic material of two parents is combined and novel genetic combinations are formed during meiotic recombination (Bell, 1982). Thus, sexual recombination is considered an important driver to establish genetic diversity (Barton and Charlesworth, 1998). However, many species, including fungi, are thought to reproduce strictly asexually (McDonald and Linde, 2002; Heitman et al., 2007; Flot et al., 2013), and have long been considered limited in their capacity to establish genetic variation. Importantly, even though asexual organisms lack meiotic recombination, adaptive evolution occurs and is established by various mechanisms ranging from single-nucleotide polymorphisms to large-scale structural variations that affect chromosomal shape, organization and gene content (Seidl and Thomma, 2014). Over longer evolutionary time-scales, all these processes establish genetic divergence that may ultimately lead to the emergence of novel species. Typically, these processes can be especially well studied in fungi that have relatively small genomes, which greatly facilitates the establishment of high-quality genome assemblies (Thomma et al., 2016).

The fungal genus *Verticillium* consists of ten soil-born asexual species with different life-styles and host ranges (Inderbitzin et al., 2011; Klosterman et al., 2011). Among these, *Verticillium* dahliae is a notorious plant pathogen that causes vascular wilt disease on hundreds of plant species, resulting in large economic losses every year (Fradin and Thomma, 2006; Klimes et al., 2015). Furthermore, also *V. longisporum*, *V. albo-atrum*, *V. alfalfae* and *V. nonalfalfae* are plant pathogens, albeit with narrower host ranges (Inderbitzin et al., 2011). The remaining species *V. tricorpus*, V. zaregamsianum, V. nubilum, *V. isaacii* and *V. klebahnii* are mostly considered saprophytes that thrive on dead organic material and that occasionally cause opportunistic infections (Ebihara et al., 2003; Inderbitzin et al., 2011; Gurung et al., 2015). Of the ten *Verticillium* species, nine are haploids while V. longisporum is an hybrid that arose from inter-specific hybridisation (Inderbitzin et al., 2011; Depotter et al., 2016). Various strains of *V. dahliae* have been sequenced (Klosterman et al., 2011; de Jonge et al., 2013), and for two strains a gapless genome assembly has been generated (Faino et al., 2015). Comparative genomics revealed the occurrence of extensive large-scale genomic rearrangements, likely mediated by erroneous double-stranded break repair, that gave rise to lineage-specific genomic regions that are enriched for in planta-expressed effector genes that encode secreted proteins that mediate host colonization (de Jonge et al., 2013; Faino et al., 2016). It is generally appreciated that effector genes are under diversifying selection pressure as the gene products are often recognized by host immune receptors as invasion patterns (Cook et al., 2015). Additionally, genomes of a single strain of *V. alfalfae* and *V. tricorpus* have been sequenced (Klosterman et al., 2011; Seidl et al., 2015).

Despite the advances in *Verticillium* genomics, the evolutionary history of this genus remains unknown. Here, we report high-quality genome assemblies of all haploid *Verticillium* species. We reconstructed ancestral *Verticillium* genomes, and reveal processes of genomic diversification that occurred during the evolution.

## RESULTS

### High-quality de novo genome assemblies of the haploid *Verticillium* species

To infer evolutionary relationships among *Verticillium* spp., we performed comparative genomics with minimum one genome of each of the nine haploid *Verticillium* species. Previously, we sequenced several *V. dahliae* strains as well as a single *V. tricorpus* strain (Klosterman et al., 2011; de Jonge et al., 2013; Faino et al., 2015; Seidl et al., 2015). Additionally, the genome of *V. alfalfae* has previously been sequenced (Klosterman et al., 2011), as well as the genomes of several *V. nonalfalfae* strains (Bioproject PRJNA283258) (Jelen et al., 2016). Additionally, we now sequenced the genomes of a strain of *V. albo-atrum V. isaacii, V. klebahnii, V. nubilum* and *V. zaregamsianum* using the Illumina HiSeq2000 platform. In total, ~18 million paired-end reads (150 bp read length of a 500 bp insert size library) and ~16 million mate-paired reads (150 bp read length of a 5 kb insert size library) were produced per strain. Subsequently, reads were de novo assembled into 35-37 Mb draft genomes (Table 1), which is comparable to the assemblies of *V. dahliae, V. tricorpus* and *V. alfalfae* (Klosterman et al., 2011; de Jonge et al., 2013; Faino et al., 2015; Seidl et al., 2015). All assemblies resulted in less than 100 scaffolds (≥ 1 kb), except for *V. isaacii* strain PD618 and *V. nubilum* strain PD621 that were assembled into 123 and 198 scaffolds, respectively (Table 1). Notably, the previously obtained assemblies of *V. tricorpus* strain MUCL9792 and *V. alfalfae* strain VaMs.102 were of significantly lower quality than the assemblies of the newly sequenced *Verticillium* species in this study (Klosterman et al., 2011; de Jonge et al., 2013; Faino et al., 2015; Seidl et al., 2015). To obtain better assemblies we decided to sequence an additional strain of each of these two species. Indeed, sequencing of *V. tricorpus* strain PD593 and *V. alfalfae* strain PD683 yielded significantly better assembly qualities, with lower numbers of scaffolds (12 for *V. tricorpus* and 22 for *V. alfalfae*) when compared with the previous assemblies (Table 1). Moreover, the assembly of *V. tricorpus* strain PD593 contained seven scaffolds with telomeric repeats at both ends, suggesting the assembly of seven complete chromosomes (Supplementary Figure S1).

**Table 1.** Assembly statistics for the various *Verticillium* genomes.

| Species | Strain name | Genome size (Mb) | # of Ns per 100_kb | N50 (Mb) | # of contigs (≥0 bp) | # of scaffolds (≥0 bp) | # of scaffolds (≥1000 bp) | CEGM A (%) | BUSCO (%) | # of proteins |
| --- | --- | --- | --- | --- | --- | --- | --- | --- | --- | --- |
| *V. dahliae* | JR2 | 36.10 | 0 | 4.2 | 8 | 8 | 8 | 95.97 | 98.75 | 10,719 |
| *V. alfalfae* | PD683 | 32.70 | 19.36 | 4.5 | 40 | 14 | 14 | 94.76 | 98.82 | 10,852 |
| *V. nonalfalfae* | TAB2 | 34.30 | 897.53 | 1.8 | 793 | 349 | 167 | 95.97 | 98.47 | 11,029 |
| *V. nubilum* | PD621 | 37.90 | 9.13 | 4.7 | 246 | 189 | 189 | 96.77 | 99.10 | 10,550 |
| *V. albo-atrum* | PD747 | 36.50 | 16.26 | 3.9 | 34 | 20 | 19 | 94.35 | 98.96 | 11,202 |
| *V. tricorpus* | PD593 | 35.00 | 14.52 | 4.4 | 71 | 11 | 9 | 95.97 | 98.82 | 10,636 |
| *V. isaacii* | PD618 | 35.80 | 62.48 | 3.1 | 239 | 188 | 122 | 95.56 | 98.96 | 10,798 |
| *V. klebahnii* | PD401 | 36.00 | 35.30 | 3.2 | 79 | 44 | 37 | 93.01 | 99.10 | 10,998 |
| *V. zaregamsianum* | PD739 | 37.10 | 55.38 | 3.5 | 75 | 46 | 32 | 94.35 | 98.96 | 11,274 |

To assess completeness of gene space, the assemblies were queried for the presence of orthologs of 248 core eukaryotic gene families using the CEGMA pipeline (Parra et al., 2007). All assemblies contained 93%-96% of these core genes (Table 1). Additionally, we also used the Benchmarking Universal Single-Copy Orthologs (BUSCO) software to assess assembly completeness with 1,438 fungal genes as queries (Simão et al., 2015). BUSCO resulted in >98% completeness for each assembly. Considering that we found CEGMA and BUSCO scores of 95.97% and 98.75%, respectively, when assessing the gapless genome assembly of *V. dahliae* strain JR2 (Faino et al., 2015), we concluded that a (near) complete gene space was assembled for all *Verticillium* species.

Next, we inferred reference gene annotations by integrating de novo and homology-based data using the Maker2 pipeline, making use of 35 predicted fungal proteomes that represent a broad phylogenetic distribution to further guide gene structure annotation (Klosterman et al., 2011; Faino et al., 2015; Seidl et al., 2015). This approach yielded around 11,000 protein-coding genes for each of the genomes, with the highest number of 11,274 genes for *V. zaregamsianum* and the lowest number of 10,636 genes for *V. tricorpus* (Table 1), which is similar to the number of genes identified in previous assemblies of *Verticillium* spp. (Faino et al., 2015; Seidl et al., 2015). However, automatic annotation of *V. dahliae* strain JR2 yielded fewer genes (10,719) than the previously generated annotation (11,430) that, next to RNA-seq data, also involved manual annotation (Faino et al., 2015). In addition to the difference in the number of predicted genes, we also observed differences in genetic features between both annotation methods, such as the overall GC% and the gene, intergenic and intron lengths (Supplementary Figure S2). However, for the coding sequences no significant differences in GC% or length were observed (Supplementary Figure S2). Thus, besides the number of predicted protein-coding genes, manual annotation mainly increased gene lengths by leveraging UTRs. As all following analyses are based on protein-coding genes, and under-estimation of gene numbers is likely similar for each of the genomes, we used Maker2 annotations for all genome assemblies.

### Phylogenetic relationships within the *Verticillium* genus

To better understand the evolutionary events during the evolution of the *Verticillium* genus, determining a robust phylogenetic relationship between *Verticillium* species is crucial. Previously, a phylogeny was constructed that was inferred from a Bayesian analysis of concatenated alignments of four protein-coding marker genes; *actin (ACT), elongation factor 1-alpha (EF), glyceraldehyde-3-phosphate dehydrogenase (GPD) and tryptophan synthase (TS)* (Inderbitzin et al., 2011). To construct the phylogenetic relationships between the nine *Verticillium* species based on whole-genome data, we used concatenated protein sequences of 5,228 single-copy orthologs and the out-group species *Sodiomyces alkalinus* to construct a maximum-likelihood phylogeny. The resulting phylogeny reveals two major clades (Figure 1), which is consistent with the previous analysis (Inderbitzin et al., 2011). The major clades are the clade Flavexudans (clade A) containing V. albo-atrum, *V. isaacii, V. klebahnii, V. tricorpus* and V. zaregamsianum, which are species producing yellow-pigmented hyphae, and the clade Flavnonexudans (clade B) containing *V. alfalfae, V. dahliae, V. nubilum* and *V. nonalfalfae*, which are species devoid of yellow-pigment.

**Figure 1.**
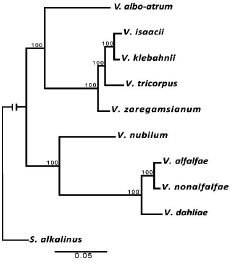
Phylogenetic tree of *Verticillium* species. Maximum-likelihood phylogeny analysis of *Verticillium* species rooted by *Sodiomyces alkalinus*. The phylogenetic tree is based on 5,228 concatenated single-copy orthologs, and the robustness of the tree was assessed using 100 bootstrap replicates.

Mitochondrial genomes have several unique characteristics, such as a conserved gene content and organization, small size, lack of extensive recombination, maternal inheritance and high mutation rates (Taanman, 1999). This makes them ideal for studying evolutionary relationships among species that diverged during a relatively short period of time. We assembled the complete mitochondrial genomes of each of the *Verticillium* species from paired-end reads using GRAbB (Brankovics et al., 2016), which extracts the reads of the mitochondrial genomes from the total pool of paired-end reads using 182 published fungal mitochondrial genome sequences as bait. We assembled all mitochondrial reads per strain into a single circular sequence containing 25 to 28 Kb with a GC content of 26-27% (Table 2), that agrees with previous assemblies for *V. dahliae* and *V. nonalfalfae* (Klosterman et al., 2011; Jelen et al., 2016). We subsequently annotated each of the mitochondrial genomes, identifying 15 protein-coding genes. For each mitochondrial genome, all genes were encoded on a single strand and in the same direction (Jelen et al., 2016) (Figure 2B). The whole-mitochondrial-genome alignments were used to construct a maximum likelihood phylogeny (Figure 2A), revealing the same topology as the phylogenetic tree that is based on the nuclear genome (Figure 1). Thus, the mitochondrial genome-based phylogenetic tree further supports the robustness of the *Verticillium* whole-genome phylogeny based on the nuclear genome assemblies, and further corroborates previous phylogenies derived by a limited set of marker genes (Inderbitzin et al., 2011).

**Table 2.** Mitochondrial genome assemblies of each *Verticillium* strain.

| **Species** | **Strain name** | **Size (bp)** | **GC content (%)** |
| --- | --- | --- | --- |
| *V. dahliae* | JR2 | 27,178 | 27.23 |
| *V. alfalfae* | PD683 | 25,167 | 26.84 |
| *V. nonalfalfae* | TAB2 | 27,203 | 26.09 |
| *V. nubilum* | PD621 | 27,203 | 27.35 |
| *V. albo-atrum* | PD747 | 28,128 | 26.58 |
| *V. tricorpus* | PD593 | 26,930 | 27.00 |
| *V. isaacii* | PD618 | 26,896 | 27.10 |
| *V. klebahnii* | PD401 | 26,871 | 27.00 |
| *V. zaregamsianum* | PD739 | 26,042 | 26.78 |

**Figure 2.**
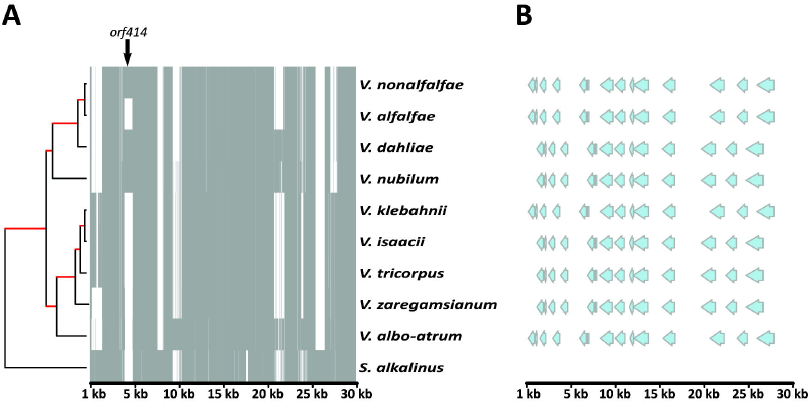
Mitochondrial genome alignments. **A** Whole mitochondrial genome alignment of all *Verticillium* species and *S. alkalinus*. Grey and white colors represent presence and absence of genomic regions, respectively. The whole mitochondrial genome alignments were used for constructing a maximum likelihood phylogenetic tree. The robustness of the topology was assessed using 100 bootstrap replicate (branches with maximum bootstrap values are in red). **B** Graphic presentation of positions of mitochondrial protein-coding genes and their orders.

It was reported recently that the mitochondrial genomes of both *V. nonalfalfae* and *V. dahliae* carry a *Verticillium*-specific region with a length of 789 bp, named *orf414* (Jelen et al., 2016). When we aligned the mitochondrial genomes of each of the *Verticillium* species and the close relative *S. alkalinus*, we found that *orf414* is absent from *S. alkalinus* (Figure 2A). However, *orf414* is not conserved in the mitochondrial genomes of all *Verticillium* species, as it is only found in *V. nonalfalfae, V. dahliae* and V. nubilum. Thus, *orf414* cannot be used as a *Verticillium*-specific diagnostic marker (Jelen et al., 2016).

### Reconstruction of ancestor genomes

The degree of synteny between extant species reveals essential information about the evolution of species from their last common ancestor (Hane et al., 2011; Lv et al., 2011). Macrosynteny occurs when large blocks of genes in the same order and orientation are shared, while microsynteny involves segments with only a handful of genes. Different from macro- or microsynteny, mesosynteny involves large blocks in which genes are conserved, but with randomized orders and orientations. In order to determine the type of synteny between the extant *Verticillium* species, we performed pairwise alignments of the genomes of the two most distantly related *Verticillium* species, *V. dahliae* and *V. albo-atrum* (Hane et al., 2011). These pairwise alignments showed clear regions of macro- and microsynteny, but no mesosynteny (Supplementary Figure S6). Subsequently, we performed similar analyses for the other *Verticillium* species, confirming the occurrence of marco- and microsynteny, and the absence of mesosynteny between *Verticillium* spp (Supplementary Figure S7).

Considering that extensive macrosynteny occurs between the extant *Verticillium* spp. we aimed to reconstruct the evolution of the *Verticillium* genus from its last common ancestor. Inferring the genome organization and gene content of ancestral species has the potential to provide detailed information about the recent evolution of descendant species. However, reconstruction of ancestral genome architectures, followed by integrating into evolutionary frameworks, has only been achieved for a limited number of species (Nakatani et al., 2007; Gordon et al., 2009; Vakirlis et al., 2016). This is largely due to either the unavailability of sequences from multiple closely related species, or due to the high fragmentation of the genome assemblies used. In order to minimize the influence of fragmented draft genome assemblies, we only considered the largest scaffolds that comprise 95% of the total set of protein-coding genes for each of the genomes (Supplementary Table S1), and constructed ancestral genome organizations that preceded each speciation event using SynChro (Drillon et al., 2014) and AnChro (Vakirlis et al., 2016). SynChro identifies conserved synteny blocks between pairwise comparisons of extant genomes, after which AnChro infers the ancestral gene order by comparing these synteny blocks. In order to validate the accuracy of AnChro, we first reconstructed the genome of the last common ancestor of *V. dahliae* and *V. alfalfae*. We did this separately for two complete and gapless genome assemblies of the extant *V. dahliae* strains JR2 and VdLs17 that, despite extensive genomic rearrangements and the presence of lineage-specific sequences (de Jonge et al., 2012; de Jonge et al., 2013; Faino et al., 2016), each contain eight chromosomes (Faino et al., 2015). Irrespective whether the genome of strain JR2 or VdLs17 was used, the resulting ancestor has nine scaffolds that are similarly organized. To compare the two ancestors, synteny blocks were constructed and aligned using SyChro, revealing an overall identical genome structure with only a few genes that lack homolog (Supplementary Figure S3). Owing to the different lineage-specific regions in the JR2 and VdLs17 genomes, the number of genes in the ancestor varies slightly, with 9,165 and 9,250 protein-coding genes based on the genome of JR2 or VdLs17, respectively. Thus, we concluded that the AnChro software is suitable for reconstruction of ancestral genomes in the genus *Verticillium*.

Ancestral genomes were reconstructed for all the nodes in the phylogenetic tree, resulting in less than 20 scaffolds at each individual node (Figure 3). Using SynChro to determine synteny blocks between the last common ancestor and the ancestor-derived genomes that served as input for ReChro (Vakirlis et al., 2016), the number of rearrangements that occurred in each branch of the tree was determined. In total, 496 rearrangements including chromosomal fusions and fissions occurred during the evolution from the last common ancestor to the nine extant haploid *Verticillium* species (Figure 3). Yet, considerable variation occurred between species ranging from 69 rearrangements for *V. tricorpus* to up to 205 for *V. albo-atrum* (Figure 3).

**Figure 3.**
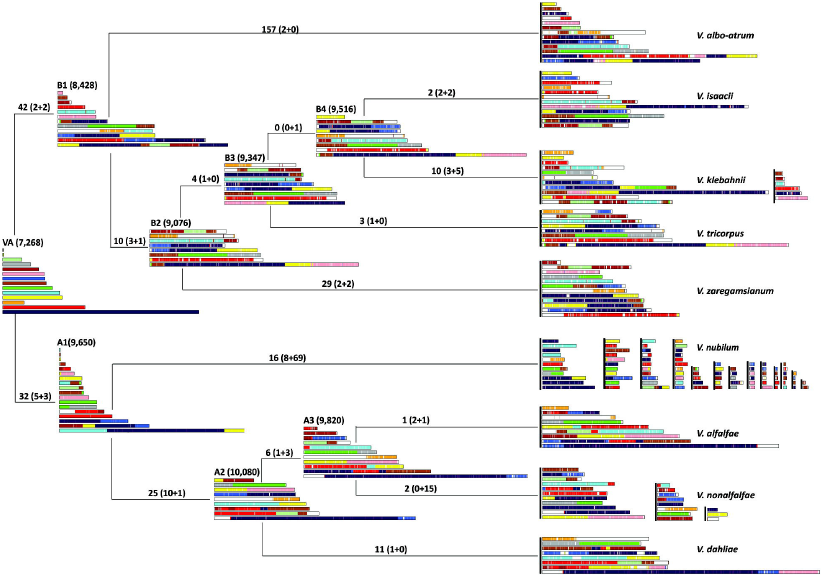
Chromosomal history of *Verticillium* genomes. Genome structure changes from the most common ancestor (VA) to the nine extant *Verticillium* species. The number of genes of each ancestors are indicted in brackets (above chromosomes). The number of chromosome sum of translocations and inversions, fusions and fissions between two genomes are indicated above each branch (the number of fusions and fissions are indicated in brackets).

To assess whether genomic rearrangements occurred in a clock-like fashion during the evolution of *Verticillium* species, we related the number of rearrangements per branch with the evolutionary time approximated by the branch lengths inferred from the phylogenetic tree (Figure 1). The branch lengths were represented by either the numbers of substitutions per site (Figure 1), or the relative divergence time that was estimated based on artificial dating of the last common ancestor of *Verticillium* and *S. alkalinus* to 100 units of time (Supplementary Figure S4). The number of rearrangements per branch showed significant correlation with both representations of branch length (R^2^= 0.3679, P= 0.007541 for substitution per site and R^2^= 0.7623, P= 6.174e-06 for relative divergence time) (Supplementary Figure S5).

### Determination of gene family expansions and contractions

To monitor gene family changes during evolution, we estimated gene family size expansion (gains) and contraction (losses) on each branch using CAFE (Han et al., 2013). CAFE models the evolution of gene family size across a species phylogeny under a birth–death model of gene gain and loss and simultaneously reconstructs ancestral gene family sizes for all internal nodes, allowing the detection of expanded or contracted families within lineages. The analysis revealed that the last common *Verticillium* ancestor contained 11,902 families with 12,631 genes. Intriguingly, *Verticillium* species generally underwent more extensive gene losses than gains (Figure 4). Among all expanded and expanded genes families, we found 1,081 gene families that are significantly more variable in size during evolution (P<0.05). The Viterbi algorithm of CAFE (P<0.05) estimates in which branches these gene families evolved more rapidly, revealing that the branches between *V. tricorpus, V. isaacii, V. klebahnii* and their last common ancestor (B4) evolved most rapidly (involving 509, 322, 242 and 165 gene families, respectively), followed by the branches of *V. alfalfae* and *V. nonalfalfae* with their last common ancestor (A3) (143 and 135 for *V. alfalfae* and A3, respectively) (Figure 4).

**Figure 4.**
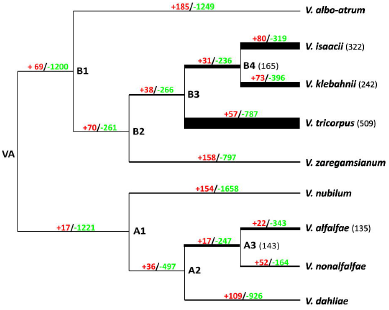
Evolution of *Verticillium* gene repertoire. The number of expanded gene families (in red) and losses (in green) were estimated on each branch of the tree under a birth–death evolutionary model. The thickened branches represent the abundance of gene families that evolved rapidly (P<0.05) and the exact number of gene families is indicated after each node name.

Next, the overrepresentation of Pfam domains in the most rapidly evolved gene families was assessed using a Hypergeometric test with a false discovery rate (FDR)-corrected P-value of <0.05. Under these conditions, only a single Pfam domain PF01636 was found to be enriched (P= 1e-06), concerning a phosphotransferase enzyme family (APH) that is predicted to consist of antibiotic resistance proteins. Potentially, this is the consequence of particular interactions between *Verticillium* species and other, antibiotic-producing, microbial species.

Although we observed considerable gene losses and gains along the different branches of the tree, we also observed a large number of shared (core) genes among all extant species. In total, 7,538 genes are common among all *Verticillium* spp., of which 1,689 are absent from *S. alkalinus* (Figure 5). Interestingly, only as little as 123 genes are clade A-specific and 288 genes clade B-specific. These genes underwent losses in the reciprocal clade in a single event only at the last common ancestor of the respective clades, rather than multiple independent losses. For each genome, less than 5% of the total protein-coding genes are species-specific (Figure 5; Supplementary data). These species-specific genes were neither enriched for any Pfam domain nor for genes that encode secreted proteins. To infer the evolutionary trajectory of these species-specific genes, we searched for homologs of these genes in the proteomes of 383 fungal species. Up to 50% of these species-specific genes had homologs in non-*Verticillium* species, suggesting that the species-specific occurrence in *Verticillium* is the result from losses from other *Verticillium* species during evolution (Supplementary Table S2).

**Figure 5.**
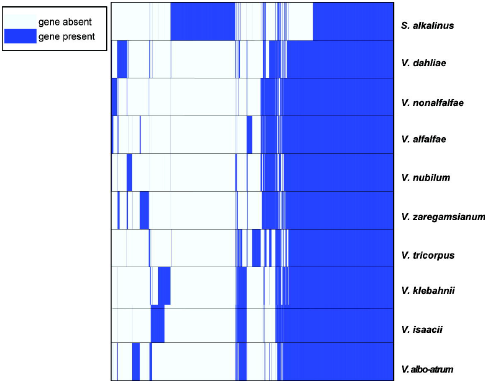
*Verticillium* gene family conservation. Presence or absence of gene families among *Verticillium* species are indicated by dark and light blue respectively. The gene families (columns) were ordered by hierarchically clustering.

To assess the characteristics of the species-specific genes versus core genes, we compared features such as GC content, gene lengths, inter-genic lengths and intron lengths. Interestingly, when compared with core genes species-specific genes significantly differ in GC content of gene sequences as well as of coding sequences, and in gene length (Figure 6A-C). Some species also show significant differences in intron lengths and intergenic lengths between species-specific and core genes (Figure 6D-E). Intriguingly, most of the species-specific genes of *V. dahliae* strain JR2 are not, or only lowly, expressed *in vitro* (Figure 6F).

**Figure 6.**
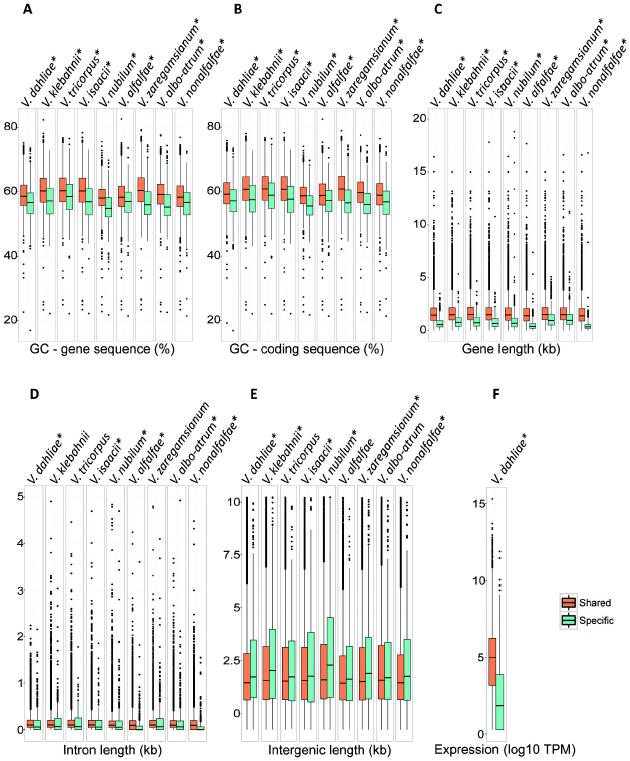
Gene feature comparisons between specific and shared genes. The Wilcoxon test (P< 0.01, indicated by asterisk) was used to detect significant differences between species-specific and shared genes of each species. **A** GC contend based on gene sequences. **B** GC contend based on coding sequences. **C** Intron length. **D** Intergenic length. **E** *V. dahliae in vitro* expression.

## DISCUSSION

Speciation occurs when two independent populations separate and get reproductively isolated. Initially the separated species will have chromosomes that share the gene content (synteny) as well as the structure and order (co-linearity). Over time, the degree of synteny and co-linearity will degrade through various processes, including chromosomal rearrangements, segmental duplications, gene losses and gene gains until, ultimately, orthologous genes in one species occur randomly in the genome of the other. In this study, we investigated genomic changes that occurred in the *Verticillium* genus from the last common ancestor to the currently recognized extant species.

We previously noted an unexpectedly high number of chromosomal rearrangements between strains of *V. dahliae* (de Jonge et al., 2013; Faino et al., 2015; Faino et al., 2016). We have speculated that the extent of rearrangements may be associated with the fact that, despite being asexual, *V. dahliae* is a successful broad host-range pathogen, reasoning that it would permit for the rapid adaptations that are required to be compatible in the arms race with host immune systems (Seidl and Thomma, 2014; Faino et al., 2016). From this hypothesis it would follow that the other species, being much less ubiquitous and successful pathogens, would not be subject to such drastic genomic rearrangements. However, our genomic reconstructions revealed that large-scale genomic rearrangements frequently occurred during speciation in the *Verticillium* genus. Moreover, our data seem to suggest that rearrangements have been even more frequent in clade A that mostly harbors non-pathogenic species. This observation makes it unlikely that the rearrangements themselves are major contributors to the pathogenicity of *V. dahliae*.

Owing to their very long evolutionary history, short generation times and abundance of asexual reproduction, the degree of evolutionary diversity within filamentous fungi is generally considered very high (Hane et al., 2011). Consequently, interspecific macro-or microsynteny is mostly lacking, even between species from the same genus (Hane et al., 2011). However, among *Verticillium* species extensive macro-as well as microsynteny is observed (Supplementary Figures 6 & 7). Previously, a particular mode of chromosomal evolution was reported to occur in particular clades of filamentous Ascomycete fungi, with genes that are conserved within homologous chromosomes but with randomized orders and orientations (Hane et al., 2011). This so-called mesosynteny was found to occur abundantly among 18 Dithidiomycete fungi (Ohm et al., 2012) but, despite being Ascomycete, not among the Sodariomycete *Verticillium* spp. (Supplementary Figures 6 & 7).

Besides extensive genomic rearrangements, conspicuous gene losses occurred during *Verticillium* evolution. This suggests that speciation is mediated by gene losses across the various lineages. Previously, gene loss was often neglected as an evolutionary driver, mostly because it was associated with the loss of redundant gene duplicates without apparent functional consequences (Olson, 1999). However, more and more genomic data suggest that gene loss acts as a manifest source of genetic change that may underlie phenotypic diversity (Albalat and Cañestro, 2016). Moreover, reduction or complete loss of gene families has been associated with ecological shifts of fungi (Casadevall, 2008). Human *Malassezia* pathogens that are phylogenetically closest related to plant pathogens such as Ustilago *maydis,* lack fatty acid synthase genes but instead produce secreted lipases to obtain fatty acids from human skin (Xu et al., 2007). Gene losses have previously also been associated with obligate biotrophic and symbiotic lifestyles of plant-associated fungi (Martin et al., 2008; Spanu et al., 2010; Duplessis et al., 2011). The impact of gene loss on speciation and pathogenicity is perhaps most evident for obligate biotrophic plant pathogens whose growth and reproduction entirely depends on living plant cells, such as the powdery mildews (Spanu et al., 2010). Their genomes show massive genome-size expansion, owing to retrotransposon proliferation, and extensive gene losses concerning enzymes of primary and secondary metabolism that are responsible for loss of autotrophy and dependence on host plants in an exclusively biotrophic life-style (Spanu et al., 2010). Recently, the evolutionary history of lineages of grass powdery mildew fungi that have a rather peculiar taxonomy with only one described species (*Blumeria graminis*) that is separated into various formae speciales (*ff.spp*.) was reconstructed. It was observed that different processes shaped the diversification of *B. graminis*, including co-evolution with the host species for some of the *ff.spp*., host jumps, host range expansions, lateral gene flow and fast radiation (Menardo et al., 2017). Arguably, as highly adapted and obligate biotrophic pathogens, powdery mildews underwent many host species–specific adaptations, which are perhaps not or less required for saprophytes and facultative and broad host range pathogens such as *Verticillium* species.

Arguably, our analyses of gene catalogues in the various species highly depend on the annotations that are generated. Previously, we manually annotated the genome of *V. dahliae* strain JR2 by making use of RNA-seq data (Faino et al., 2015). As the comparison of genomes that are annotated with different methods would introduce artefacts in our findings, RNA-seq data are not available for each of the species and manual annotation of every genome is not feasible, we automatically annotated all the genomes that were used in this study. Based on the comparison of the manual and automatic annotation of the genome of *V. dahliae* strain JR2, we likely under-estimate the gene content of the various genomes, but arguably to the same extent for the various genomes. We therefore anticipate that the manual annotation in itself does not impact the major findings of this study.

It is tempting to speculate that effector catalogs, especially those parts that are lineage-specific inventions, arise by gene acquisition often through gene duplication followed by neo-and subfunctionalization. Alternatively, genes can be acquired via horizontal gene transfer. We have previously shown that the important virulence effector Ave1 that mediates aggressiveness of *V. dahliae* on various host plants was acquired via horizontal gene transfer (de Jonge et al., 2012). Moreover, extensive study of *V. dahliae* genomes revealed that lineage-specific regions that harbor in planta-expressed effector genes arose by segmental duplications, likely generating the genetic material that subsequently has the freedom to diverge into novel functions (de Jonge et al., 2013; Faino et al., 2016).

Functions of species-specific genes are often found to be associated to species-specific adaptations to a certain environment (Domazet-Loso and Tautz, 2003). For example, morphological and innate immune differences among *Hydra* spp. are controlled by species-specific genes [52, 53]. In plant pathogens, many effectors characterized so far are species-specific and facilitate virulence on a particular host plant [54]. However, these species-specific effectors are frequently mutated, or even purged, in order to overcome host recognition (Cook et al., 2015). Moreover, it has been shown that compared to conserved genes, species-specific genes are frequently shorter (Lipman et al., 2002), evolving quicker and less expressed (Domazet-Loso and Tautz, 2003; Plissonneau et al., 2016). Consistent with this observation, our results show that species-specific genes differ in their characteristics compared to core genes. It has often been claimed that these distinct characteristics are hallmarks of gene structure degeneration and that these species-specific genes are more likely to go extinct (Palmieri et al., 2014; Plissonneau et al., 2016). Thus, even though species-specific genes in *Verticillium* may have important functions, they may be more likely to get lost during evolution.

## EXPERIMENTAL PROCEDURES

### Genome sequencing and assembly

DNA was isolated from mycelium of 3-day-old cultures grown in potato dextrose broth (PDB) at 28°C as described previously (Seidl et al., 2015). Of each strain, two libraries (500 bp and 5 Kb insert size) were sequenced using the Illumina High-throughput sequencing platform (KeyGene N.V., Wageningen, The Netherlands). Genome assemblies were performed using the A5 pipeline (Tritt et al., 2012), and sequence gaps were filled using SOAP GapCloser (Luo et al., 2012). Next, QUAST (Gurevich et al., 2013) was used to calculate genome statistics. Illumina sequence reads and assemblies were deposited in NCBI (Bioproject PRJNA392396).

### Gene prediction and annotation

Protein-coding genes were de novo annotated with the Maker2 pipeline (Holt and Yandell, 2011) using 35 predicted fungal proteomes and the previously annotated proteomes of *V. dahliae* and *V. tricorpus* to guide gene structure annotation (Faino et al., 2015; Seidl et al., 2015). Secretome prediction was described previously (Seidl et al., 2015).

### Ortholog analysis and tree building

Ortholog groups were determined using OrthoMCL (Li et al., 2003). The species phylogenetic tree was generated using 5,228 single-copy orthologs that are conserved among all of the genomes. Individual families were aligned using mafft (LINSi; v7.04b) (Katoh et al., 2002) and concatenated. Maximum likelihood phylogeny was inferred using RAxML (v7.6.3) (Stamatakis, 2014). The robustness of the inferred phylogeny was assessed by 100 rapid bootstrap approximations.

### Mitochondrial genome assembly and comparison

Sequencing reads of mitochondrial genomes of each *Verticillium* species were extracted from raw paired-end reads using GRAbB using 182 already published fungal mitochondrial genome sequences as bait (Brankovics et al., 2016). Subsequently, GRAbB assembled all the mitochondrial reads into single circular sequences. The mitochondrial protein-coding genes were determined by using already published *Verticillium* dahliae protein sequences BLAST against mitochondrial genomes. Whole mitochondrial genome alignment was performed by mafft (default setting) (Katoh et al., 2002), and the likelihood phylogenetic tree was built using RAXML (v7.6.3) (Stamatakis, 2014). The robustness of the inferred phylogeny was assessed by 100 rapid bootstrap approximations.

### Ancestor genome reconstruction and comparison

Ancestral genomes were constructed by using CHROnicle package that comprises SynChro (Drillon et al., 2014), ReChro and Anchro (Vakirlis et al., 2016). Conserved synteny blocks were identified between pairwise combinations of genomes with SynChro (Drillon et al., 2014). The synteny block stringency delta, which determines the maximum number of intervening Reciprocal Best Hits (RBH) allowed between anchors within a synteny block was set to three. Subsequently, ReChro was used for estimating genome rearrangements from ancestral to extant genomes.

### Gene family gains and losses

Gene family gain/loss analysis was carried out using CAFE (Han et al., 2013). Pfam function domains were predicted using InterProScan (Jones et al., 2014). For analyses of Pfam domain associated with gene families (OrthoMCL clusters), Pfam domains were assigned to specific families (clusters) only when present in at least half of the homologs within each gene family. Pfam enrichment of gene families of interests was carried out using hypergeometric tests, and significance values were corrected using the Benjamini-Hochberg false discovery method (Benjamini and Hochberg, 1995).

## ACKNOWLEDGEMENTS

We thank Drs Inderbizin and Subbarao for sharing fungal isolates and Sander Rodenburg for help with scripting.

---

Supplementary Figure S1. Overview of the draft genome assembly of *V. tricorpus*, strain PD593. Schematic representation of the eight largest scaffolds in the genome assembly of *V. tricorpus* strain PD593. Characteristic fungal telomeric repeats are displayed on the ends of the scaffolds (indicated by red color).

Supplementary Figure S2. Gene feature comparisons between automatically and manually annotated genes of V dahliae strain JR2. **A** GC content. **B** Gene/coding sequence length. **C** Intergenic length. **D** Intron length.

Supplementary Figure S3. Chromosome alignments between two ancestor genomes. Two ancestor genomes were reconstructed based on the genome of two *V. dahliae* strain JR2 and VdLs17 respectively (indicated by red and blue dots respectively). Green and red plain lines highlight homology relationships with same and inverted directions, respectively.

Supplementary Figure S4. Ultrametric phylogeny of *Verticillium* species and *S. alkalinus*. The ultrametric tree derived by a maximum likelihood analysis of concatenated single-copy orthologs (Figure 1). Averaged divergence times per branch are reported relative to an arbitrary age of the last common ancestor of *Verticillium* spp. and *S. alkalinus* (set at 100).

Supplementary Figure S5. Correlations between the branch lengths and number of rearrangements. **A** Branch length is represented by number of substitutions per site (Figure 1), **B** Branch length is represented by evolutionary distance (Supplementary Figure 3)

Supplementary Figure S6. Dot-plot comparison between *V. dahliae* and V. albo-atrum. The six-frame translations of both genomes were compared using Promer (Mummer 3.0). Homologous regions are plotted as dots and are color coded for percentage similarity. Macro-or micro-synteny is indicated by long or short diagonal lines with a slope that is positive or negative depending on whether the genes align in the same or inverted order.

Supplementary Figure S7. **A-D** Pairwise Dot-plot comparison between nine haploid *Verticillium* species. Macro-or micro-synteny is indicated by long or short diagonal lines with a slope that is positive or negative depending on whether the genes align in the same or inverted order.

